# Social Connection: Resting-State Functional Connectivity Associations with Social Networks, and the Roles of Empathy and Sex

**DOI:** 10.64898/2026.11.12.766105

**Authors:** Mia G Casburn, Andrea Fernandes, Lara Ressin, Alan Sheerin, Theodora Voulgari, Aodán Laighneach, Derek W Morris, Dara M Cannon

**Affiliations:** Clinical Neuroimaging Laboratory, Center for Neuroimaging, Cognition and Genomics (NICOG), Galway Neuroscience Center, School of Medicine, College of Medicine, Nursing and Health Sciences, University of Galway, Galway, Ireland; School of Biological and Chemical Sciences and Institute for Health Discovery and Innovation, University of Galway, Galway, Ireland

**Author notes:** Joint senior authors (Derek W Morris, Dara M Cannon).

**Keywords:** social network index, empathy, power spectra, default mode network, independent component analysis, salience network, social brain

## Abstract

Smaller social networks (SNs) negatively influence cognition, psychiatric trajectories and mortality. Studies linking SNs to resting-state functional connectivity (FC) inconsistently suggest orbito-frontal and frontoparietal regions and implicate a potential role for empathy and sex. We investigate how SN characteristics relate to FC, additionally assessing if empathy and sex alter these associations.

Group-independent component analysis was applied to multi-band resting-state fMRI from 83 participants (Caltech Conte dataset: ds003798). The Social Network Index measures SN-contacts (SN-contacts) SN-diversity (number of roles) and SN-embeddedness (number of contact groups) and the Empathy Quotient measured empathy. Using MANCOVA, we investigate if resting-state network power and connectivity are associated with SNI measures and whether these relationships differ by empathizing ability and sex.

Participants aged 18-48 (mean age 29.3±7.0, 55% male) demonstrated SN-contacts and SN-diversity associated with lower frontal-pole FC within the default-mode network (DMN, q-values<0.05). Empathy altered how SN-diversity and SN-contacts relate to superior lateral-occipital, default mode and salience network FC (q-values<0.05). Finally, males and females showed diverging associations of SN-contacts and SN-diversity with salience network power.

Smaller SNs were associated with increased anterior-DMN FC, potentially indicating a tendency towards internally-directed cognition in more socially isolated individuals. Opposing relationships between SNs with salience and lateral-frontoparietal networks by empathy level highlights empathy as critical for understanding social biology. Males and females differed in SN associations with salience network coherency, supporting known sex-specific neural correlates of social behavior. These findings provide insights into the biology of social networks, offering potential targets for studying social deficits in psychiatric populations.

## 1. Introduction

The social brain hypothesis originated in studying primates, theorizing that inherently social species developed larger, more efficient brains to adjust to the cognitive demands of social life and assist species survival (Dunbar, 1998; Fishman et al., 2025). Extended to humans, this hypothesis proposes that individuals who are more socially active have increased gray matter in specific brain regions termed the “social brain”, offering a potential explanation for individual variability in social tendencies (Bickart et al., 2011; Oesch, 2024). Additionally, smaller social networks and social isolation are linked with poorer health, including increased mortality risk (Cacioppo & Cacioppo, 2014; Holt-Lunstad et al., 2010) higher depression (Fiori et al., 2006; Santini et al., 2020) cognitive impairment (Cacioppo & Hawkley, 2009; B. Kim et al., 2026) and more severe psychiatric symptoms in patient populations (Degnan et al., 2018). Both the social brain hypothesis and social health research provide a framework for investigating associations between social behavior and biology to better understand individual differences in social behavior, social deficits in psychiatric disorders, and the overall impact of these on health outcomes (Dunbar, 2025; Fishman et al., 2025). Resting-state functional MRI (rsfMRI) indirectly measures brain activity through fluctuations in blood-oxygen levels in the brain, reflecting changes in local field potentials representing synaptic inputs and their subsequent processing within a population of neurons (Logothetis, 2003; Logothetis et al., 2001) with low-frequency brain fluctuations at rest reflecting meaningful neural organization (Allen et al., 2011; Biswal et al., 1995). This offers the potential to examine how dynamic brain networks may relate to social behavior and health.

In line with the social brain hypothesis, previous region-of-interest (ROI) studies have identified associations between resting-state functional connectivity (FC) networks and social network measures. For example, Joo *et al*. (2017) found associations between social networks and FC between the right inferior frontal/insular cortices and the superior and middle occipital gyri. Pruitt *et al*. (2024) showed that salience network (anterior cingulate and insula) connectivity was positively associated with social engagement and mediated the association between social engagement and cognition. Others implicated amygdala-orbitofrontal cortex FC (Bickart et al., 2012; Noonan et al., 2018; Zou et al., 2016). Although important for identifying candidate regions associated with social behavior, these hypothesis-driven approaches may bias findings or overlook network-level effects. Alternatively, data-driven whole-brain studies (Casburn et al., 2026; Lin et al., 2020) can offer insights into how different regions dynamically co-ordinate activity across the brain to promote social networks. Accordingly, Pillemer *et al*. (2017) found within-network FC of the sensorimotor, visual and frontoparietal networks associated with social networks while Kiekhaefer *et al*. (2023) found similar results when investigating FC associations with social activity participation, also implicating frontoparietal and somatomotor networks.

Evidently, findings on which brain phenotypes are associated with social network characteristics remain inconsistent, with the main finding corroborated across studies implicating the insula (Joo et al., 2017; Pillemer et al., 2017; Pruitt et al., 2024) although this is ascribed to different brain networks across studies. The lack of whole-brain investigations is just one of many factors likely contributing to this variability. Additional limitations include investigations into FC and social network measures largely consisting of small sample sizes (n=19 to 61) (Bang et al., 2019; Noonan et al., 2018; Pillemer et al., 2017; Zou et al., 2016) limiting statistical sensitivity. Many studies also tend to exclusively focus on either younger (Bickart et al., 2012; Mori & Haruno, 2022) or older (Joo et al., 2017; Pillemer et al., 2017; Pruitt et al., 2024) participants with few including middle-aged individuals. Existing moderately sized studies (Mori & Haruno, 2022; Pruitt et al., 2024) use region-based methods, which may bias findings or overlook network-level influences. Some investigations do utilize larger samples such as the UK Biobank (Anatürk et al., 2021; Casburn et al., 2026; Kieckhaefer et al., 2023) however these are limited by shallow social phenotyping and primarily retired participants, further complicating comparability across findings. Together, these limitations highlight the inhomogeneity and limited generalizability in existing studies investigating functional connectivity associations with social phenotypes.

One of the most commonly used social phenotypes across studies is the Social Network Index (SNI) a measure of multiple aspects of real-world social networks consisting of subscales indexing network complexity, diversity and size (Cohen et al., 1997). Existing studies linking the SNI to brain connectivity tend to exclusively use one of the three SNI sub-scales, but research suggests that distinct characteristics of social networks may provide unique social resources and therefore differentially relate to various outcomes such as mental health (Fiori et al., 2006) and brain phenotypes (Liu et al., 2018). Consequently, the core objective of this study is to investigate the functional brain associations of the three SNI sub-scales to establish whether they are characterized by distinct or overlapping neural correlates using a data-driven approach.

Extending this, considering a persons’ social network independently, as well as in relation to their social-cognitive ability, may prove more conceptually informative. Kennedy & Adolphs (2012) describe social behavior as linked to the brain via social cognition, where brain structure and function facilitate cognitive processes that enable social behavior. Empathy is a core social ability that can be composed of cognitive and affective elements (Baron-Cohen & Wheelwright, 2004; Oliveira Silva et al., 2018) that allow humans to understand the intentions and mental states of others, predict their behavior, and feel in response to another person’s emotional state. Existing studies have found that cognitive empathy mediates the association between the SNI with FA values in the cingulum (Veerareddy et al., 2025) and volume of the dorso-lateral pre-frontal cortex (Veerareddy et al., 2023). Powell *et al*. (2012) also found that mentalizing/intentionality, a social-cognitive ability sharing neural correlates with empathy (Arioli et al., 2021) mediated the association between orbitofrontal volume and social network size. These findings suggest that empathizing may provide an explanatory mechanism linking structural brain characteristics to social networks.

While these structural studies provide valuable insight, social behaviors and cognition are likely supported by dynamic communication between distributed brain regions rather than isolated structures (Kennedy & Adolphs, 2012). Therefore, functional brain measures may provide a novel insight into how communication between brain regions relates to empathic response and social behavior. This relationship has yet to be investigated in the context of functional brain measures, presenting a significant gap in the literature. Cognitive empathy has been linked to resting-state FC measures, including within default mode (Winters et al., 2021) sensorimotor, salience, and central executive regions (Bilevicius et al., 2018). Moreover, both cognitive and affective empathy have been positively associated with social network size (Kardos et al., 2017) and social network embeddedness (Wölfer et al., 2012) making empathy a plausible moderator for the relationship between FC and SNI measures. Accordingly, a secondary objective of this study is to establish whether empathy interacts with SNI, querying if the association between resting-state networks and the SNI subscales differs with varying empathy scores. Instead of focusing on a specific sub-element of empathy, we utilize the Empathy Quotient, a self-report measure of global empathy reliable across both clinical and healthy samples (Baron-Cohen & Wheelwright, 2004; Lawrence et al., 2004). Investigating this relationship not only provides a cohesive insight into social brain and behavioral links but may offer a potential target for understanding social deficits in psychiatric disorders, via empathetic ability. Therefore, we propose a model where the relationship between resting-state network connectivity and real-world social behavior may depend on differences in empathizing abilities (Kennedy & Adolphs, 2012).

With a goal to establish factors that may alter SNI and FC relations, an additional final aim of this study is to investigate whether this association differs in males and females. The relationship between brain structure and social behavior differs across sexes (Kiesow et al., 2020) with prominent sex differences observed in general functional connectivity (Allen et al., 2011) and EEG responses to socially valanced stimuli (Proverbio et al., 2008). Additionally, males and females differ in the size and characteristics of their social groups (David-Barrett et al., 2015; Dunbar, 2025; Dunbar et al., 2024) and social-cognitive abilities (Proverbio, 2017, 2023). However, our understanding of sex differences in the social brain is limited (Pavlova, 2017) making the interaction between SNI and sex on FC measures a relevant factor to consider.

Overall, current evidence suggests a relationship between brain connectivity and social networks, however, whether this relationship is influenced by empathizing ability and sex remains largely understudied. The current study uses group independent-component-analysis (GICA) (V. D. Calhoun et al., 2001) to examine the association between resting-state networks and social network characteristics, and investigates whether this relationship varies across birth sex and individual differences in empathy. Given that existing studies linking social phenotypes to brain connectivity tend to implicate widespread brain regions across a variety of different networks (Kennedy & Adolphs, 2012) an ICA-based approach allows for a data-driven method of establishing wide-scale brain networks associated with the SNI without constraining findings by pre-defined regions-of-interest. To overcome existing limitations, we combine this data-driven approach with a moderately sized participant sample (n = 83, Kliemann et al., 2022) and utilize all three SNI metrics. Outcome measures include both spatial maps indexing the functional connectivity and spatial topology of each network, and power spectra, representing network coherency as the power of amplitude fluctuations across frequencies (Allen et al., 2011). The aims are twofold: firstly, to establish relationships between social network characteristics with rsfMRI spatial maps of connectivity and power spectra, and secondly, to investigate whether empathy and/or sex interact with social network characteristics, changing how this relationship manifests. Although *a priori* hypotheses regarding specific networks were not defined, we anticipate that networks previously reported as supporting social cognition, such as the default mode network, will be associated with the SNI, and that these relationships will differ depending on sex and EQ scores.

## 2. Methods

### 2.1 Study Design & Participants

The data used in this study was derived from an open-access dataset available on OpenNeuro acquired by the Caltech Conte Center lab at the California Institute for Technology (https://openneuro.org/datasets/ds003798/versions/1.0.5). Kliemann *et al*. (2022) provide detailed information about the dataset, image acquisition, processing steps, and quality control. Data was collected pertaining to 191 adults between the ages of 18-50 with a final sample of 117 due to exclusion criteria (Table 1) and participant drop-out. We further excluded participants due to mixed acquisition protocols, including only individuals with multi-band imaging and all SNI measures, resulting in a final sample of 83.

**Table 1.** Participant Exclusion Criteria

| Category |  | Criteria |  |  |
|---|---|---|---|---|
| Current | – | Full-Scale IQ < 90 |  |  |
|  | – | English non-fluency |  |  |
|  | – | Uncorrected vision/hearing impairment |  |  |
|  | – | First-degree relative with schizophrenia/autism spectrum |  |  |
|  | – | Beck Depression Inventory-II Score > 25 |  |  |
|  | – | Psychotropic medication use |  |  |
| Medical History | – | Premature birth | – | Metabolic disorder |
|  | – | Epilepsy | – | Chemotherapy/radiation |
|  | – | Head Injury | – | Brain surgery |
|  | – | Eating Disorder | – | Psychosis |
|  | – | Neurological condition | – | Bipolar Disorder |
|  | – | Previous suicide attempt | – | Autism |
|  | – | Substance dependence/abuse | – | Alcoholism |
|  | – | Color blindness |  |  |
| MRI-Acquisition Related | – | Claustrophobia | – | Pregnancy |
|  | – | Incompatible tattoos | – | Structural abnormality detected |
|  | – | Excessive motion during scan | – | MRI incompatibilities |
| Data-Related | – | Single-band imaging | – | No SNI or EQ data |

### 2.2 Social & Empathy Phenotypes

The Social Network Index (SNI) is a self-report questionnaire assessing social networks across 12 social roles including parent, spouse, child, close relative, child-in-law, neighbor, close friend, church member, employee, student, group member and volunteer (Cohen, 1991) and includes three sub-scales. Number of social contacts measures the number of people the individual has contact with at least once every 2 weeks. It showed a positively skewed distribution (skew = 1.42, kurtosis = 2.69) and was log transformed (log(SN contacts +1)) resulting in an improved distribution (skew = -0.96) and reduced kurtosis (1.44). Social network diversity measures the number of social roles (e.g. close friend, parent, etc.) where an individual has contact with one person at least once every 2 weeks with a maximum of 12 and was normally distributed (skew = -0.27, kurtosis = 0.07). Finally, social network embeddedness measures the number of different social groups/domains (e.g. family, friends, work, etc.) these contacts belong to. A point is given if the individual has at least 4 contacts in that domain (e.g. 4 work contacts and 4 friend contacts give a score of 2) with a maximum of 8. SN embeddedness was right skewed with very few participants scoring greater than three (n = 7) therefore it was converted into a binary variable with individuals reporting 0 or 1 social domains scored as “Limited Social Embeddedness” and those reporting 2 or more as “Socially Embedded”.

Empathy was measured using the Empathy Quotient (Baron-Cohen & Wheelwright, 2004; Lawrence et al., 2004) an established 40-item self-report scale encompassing cognitive and affective empathy. It is an additive score measured through 40 questions, half of which require an agreement to add to the score and half require a disagreement, where a “definite” is scored with a 2 and a “slight” is scored with a 1, with a maximum score of 80 (Table 2).

**Table 2.** Example of scoring system for Empathy Quotient

| **Prompt** | **Definitely Disagree** | **Slightly Disagree** | **Slightly Agree** | **Definitely Agree** |
|---|---|---|---|---|
| “I enjoy caring for other people” | 0 | 0 | 1 | 2 |
| “Seeing people cry doesn’t really upset me” | 2 | 1 | 0 | 0 |

### 2.3 Image Acquisition

Rs-fMRI scans (3T) were acquired in two runs of 420 seconds using multi-band isotropic T2*-weighted EPI, depending on protocol versions which changed throughout recruitment (Kliemann et al., 2022). Images were acquired with two different protocols, both consisting of 2.5 mm isotropic voxels and 30ms TE but differing in TR and flip angle (multi-band protocol 1: 1000ms TR and flip angle 60 degrees; multi-band protocol 2: 700ms TR and flip angle 53 degrees).

### 2.4 Image Processing

Minimally pre-processed functional images were obtained from the open-access dataset provided on OpenNeuro. These had already been pre-processed using the fMRIprep 20.2.1 pipeline (Esteban et al., 2019) and a full fMRIprep description is provided in Supplementary Box 1. Briefly, T1-weighted images used for registration were corrected for intensity inhomogeneities, skull-stripped and segmented into CSF, white-and gray-matter using ANTs 2.3.3 (Avants et al., 2009) Freesurfer 6.0.1 (Fischl, 2012) and FSL 5.0.9 (Jenkinson et al., 2012) and non-linearly registered to the MNI152NLin2009cAsym template using ANTs.

Rs-fMRI pre-processing included generation of a reference BOLD volume and corresponding skull-stripped image for all runs by aligning and averaging single-band reference images. BOLD images were susceptibility distortion corrected and co-registered to each subject’s T1-weighted image using boundary-based registration. Head-motion parameters were estimated using FSL MCFLIRT (Jenkinson et al., 2002). All spatial transformations were applied in a single resampling step, and preprocessed BOLD images were normalized to the MNI152NLin2009cAsym standard space.

The fMRIprep outputs were then subject to further processing. Multi-band images contained initial non-steady state volumes as identified by fMRIprep: therefore, the first ten volumes were removed. Although ICA networks can be identified without spatial smoothing, low FWHM values may introduce additional anti-correlations (Alahmadi, 2021). Therefore, a medium smoothing kernel (FWHM 5mm) was selected as the most appropriate (Candemir, 2023; Chen & Calhoun, 2018) also aligning with recommendations to use approximately double the voxel size (Kumar et al., 2024; Pajula & Tohka, 2014; Worsley & Friston, 1995). A corresponding Gaussian sigma value was calculated based on the formula 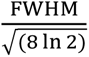 giving σ = 2.123mm which was applied using fslmaths -s.

### 2.5 Group Independent Component Analysis

Prior to ICA, the optimal number of components, 42, was determined using the Minimum Description Length (MDL) i.i.d criteria (Y. Li et al., 2007). GICA is a blind-separation technique that starts with subject-level data reduction using principal components analysis (PCA) then temporally concatenates across subjects and further reduces using a group-level PCA step, ultimately extracting independent components (ICs) from group level data (V. D. Calhoun et al., 2001). We implemented a group-level ICA to derive 42 ICs within the GIFT toolbox (V. D. Calhoun et al., 2001, 2009) using the Infomax algorithm, as it has been shown to exhibit higher reliability across runs than other non-deterministic ICA algorithms when applied to fMRI data (V. Calhoun et al., 2004; Correa et al., 2007; Erhardt et al., 2011; Esposito et al., 2002; Wei et al., 2022). InfoMax maximizes the entropy of output components, minimizing mutual information to derive statistically independent components (Bell & Sejnowski, 1995).

Finally, we derived subject-specific connectivity spatial maps and time courses using the Group Information Guided (GIG-ICA) method (Du & Fan, 2013). This uses a multi-objective optimization approach balancing individual variability of subject-level components with the matching of components to the group-model, is more sensitive to individual differences (Salman et al., 2019) and derives more reliable components across subjects than other back reconstruction methods (Du et al., 2016). Following ICA, component stability was assessed with ICASSO which selects an optimal number of stable components across repeated ICA runs, excluding components with an Iq value < 0.8.

### 2.6 Resting-State Network Classification

Components with an Iq value > 0.8 were then classified as either noise or signal adhering to the guidelines provided by Griffanti *et al*. (2017). This involved an independent rating procedure where two raters (MC, LR) established whether a component was signal based on it consisting of high-power in predominantly low-frequencies (0.01-0.1 Hz) with peak activations and large clusters located in the grey matter (Griffanti et al., 2017). Cohen’s kappa was estimated to test inter-rater reliability, and any disagreements were resolved by two additional raters (AF, DMC).

Finally, the components deemed to represent signal were then independently assessed for correspondence with known resting state networks (Beckmann et al., 2005; Laird et al., 2011; Shirer et al., 2012; Smith et al., 2009; Uddin et al., 2019) again by the same two independent raters (MC, LR) with Cohen’s kappa calculated and other raters consulted if required (AF, DMC).

### 2.7 Outcome Measures

Outcome measures included both spatial maps of connectivity and power spectra for each component. Spatial maps index the spatial coactivation/functional characteristics of each component while power spectra characterize frequency-specific amplitudes of spontaneous network fluctuations, providing insights into the temporal dynamics of intrinsic network activity (Allen et al., 2011). Power spectra were normalized by the total spectral power and log-transformed, representing proportion of the power at each frequency. Spectra were examined up to a cut-off of 0.2 Hz, extending the traditional higher cut-off of 0.1Hz based on studies showing that higher frequencies also contribute to the BOLD signal (Baria et al., 2011; Niazy et al., 2011).

### 2.8 Statistical Analyses

To identify associations between the SNI scales and functional connectivity component features (spatial maps and power spectra) we used the MANCOVAN Toolbox (Allen et al., 2011; Gruner, 2026). This toolbox uses a backward selection procedure where all effects are initially tested at a multivariate level (i.e. the average spatial map loading of the whole component associated with SNI) if this is significant this is then carried through to the univariate level (i.e. individual component voxels associated with SNI). All univariate results were corrected for multiple comparisons using false discovery rate q-values (q<0.05). The SNI measures were significantly and positively associated with each other (Table S1, r range = 0.64-0.86, all ps < 0.001) therefore necessitating the use of individual models. This resulted in three models including main effects of SNI, EQ, age and sex and interactions of each SNI variable with EQ and sex, controlling for mean framewise displacement (FWD) and acquisition variability as nuisance regressors.

To assess the robustness of these findings, a leave-one-out sensitivity analysis was conducted. For each effect the analysis was run 83 times, removing a different participant on each iteration. An effect was considered robust if it remained significant (p < 0.05) at each iteration. All results reported passed this sensitivity check (Tables S4 and S5) unless otherwise specified.

## 3. Results

### 3.1. Participants

Participants (n = 83) were 55% male with a mean age of 29 (range 18-49, Table 3). EQ was positively associated with SN contacts (*r*(81) = 0.28, *p* = 0.009) and SN diversity (*r*(81) = 0.28, *p* = 0.009) but not social embeddedness (r(81) = 0.19, p = 0.08, Table S1). Females had higher mean SN diversity (mean = 5.11 ± 1.65) compared to males (mean = 4.26 ± 1.82, W = 627.50, p = 0.04) and significantly higher mean EQ scores (mean = 53.30 ± 11.23) compared to males (mean = 46.21 ± 12.21, W = 550, p = 0.006). None of the SNI variables or EQ exhibited significant age associations and also didn’t differ across different races or education levels (Table S2).

**Table 3.** Sample Demographic Characteristics

| **Demographic** | **Frequency** | **Percentage** |
|---|---|---|
| **Sex at Birth** |  |  |
| Male | 46 | 55% |
| Female | 37 | 45% |
| **Age (mean 29.29 ±6.96)** |  |  |
| 18-22 | 11 | 13% |
| 23-27 | 31 | 37% |
| 28-32 | 17 | 21% |
| 33-37 | 10 | 12% |
| 38-42 | 10 | 12% |
| 43+ | 4 | 5% |
| **Race** |  |  |
| African American/Black | 8 | 10% |
| Asian | 12 | 14% |
| Caucasian/White | 37 | 45% |
| Multiracial | 17 | 20% |
| Other | 9 | 11% |
| **Highest Education Level** |  |  |
| High School | 4 | 5% |
| Some College | 25 | 30% |
| Associate’s degree | 10 | 12% |
| Bachelor’s Degree | 37 | 45% |
| Postgraduate Degree | 7 | 8% |

### 3.2. Resting-State Networks Classification

Of the 42 components derived from the ICA, 3 were deemed unreliable, leaving 39 components for signal or noise classification. Independent raters (MC & LR) initially showed moderate inter-rater reliability (Cohen’s κ = 0.56) following consultation with two additional raters (DMC & AF) final consensus identified 16 components (41%) as noise with the remaining 23 (59%) retained for subsequent analyses.

Neuroanatomical classification into established resting-state networks exhibited an inter-rater reliability indicating substantial agreement (Cohen’s κ = 0.70) and resulted in component groups of lateral frontoparietal (two right lateralized, one left, and one bilateral) default mode (four) salience (three) sensorimotor (three) lateral visual (two) language (one) auditory (one) cerebellar (one) pre-frontal (one) default mode/salience combination (one) lateral frontoparietal/language combination (one) and medial visual (one) (Table 4).

**Table 4.** Independent-Component Analysis-Derived components representing Resting-State Networks

| Component | Neuroanatomy | MNI Co-ordinates | Assigned network |
|---|---|---|---|
| Component 2 | 54% L pre-central, 32% post-central<br>44% R pre-central, 24% post-central | -60, -4, 26<br>60, -2, 24 | Sensorimotor |
| Component 8 | 31% L post-central, 21% pre-central<br>50% R post-central, 23% supramarginal<br>61% L supplementary motor (juxtapositional lobule) | -36, -25, 71<br>50, -28, 52<br>-2, -12, 52 | Sensorimotor |
| Component 30 | 28% R post-central, 8% posterior supramarginal<br>42% R post-central, 18% pre-central<br>83% R inferior lateral occipital | 48, -34, 62<br>48, -14, 16<br>50, -68, 0 | Right Sensorimotor |
| Component 20 | 77% L frontal pole<br>31% R frontal pole, 28% middle frontal | -33, 59, 14<br>36, 38, 24 | Pre-frontal |
| Component 36 | 72% L superior lateral occipital<br>52% L precuneus, 11% supracalcarine, 6% intracalcarine<br>53% R precuneus, 23% supracalcarine | -39, -82, 38<br>-12, -58, 14<br>18, -58, 16 | Default Mode |
| Component 29 | 22% R frontal pole, 5% superior frontal<br>86% R frontal pole<br>61% R paracingulate, 10% superior frontal | 0, 59, 26<br>26, 60, 18<br>8, 48, 20 | Anterior Default Mode |
| Component 18 | 73% R angular, 10% middle temporal (temporooccipital)<br>55% inferior frontal (pars triangularis) 19% frontal pole<br>47% precuneus, 35% posterior cingulate<br>24% middle temporal anterior, 20% middle temporal posterior, 16% superior temporal anterior, 14% superior temporal posterior | 60, -52, 17<br>56, 32, 2<br>8, -50, 36<br>58, -8, -12 | Default Mode |
| Component 11 | 68% precuneus, 18% cuneal cortex<br>48% posterior cingulate | 0, -76, 38<br>6, - 28, 28 | Posterior Default Mode |
| Component 37 | 26% superior frontal, 15% paracingulate<br>37% middle frontal, 19% inferior frontal (opercularis)<br>43% frontal orbital, 23% frontal operculum | 0, 41, 38<br>52, 16, 34<br>44, 26, -6 | Default Mode-Salience |
| Component 22 | 41% medial frontal pole<br>59% L paracingulate, 37% anterior cingulate<br>28% frontal operculum, 7% frontal orbital, 7% temporal pole, 5% inferior frontal (opercularis) 5% insula | 0, 62, 8<br>-4, 44, 16<br>48, 16, -4 | Salience |
| Component 24 | 38% R inferior frontal (opercularis) 8% precentral<br>35% L temporal pole, 17% inferior frontal (opercularis)<br>34% R paracingulate, 23% supplementary motor (juxtapositional lobule) | 57, 14, 1<br>-56, 8, -2<br>2, 80, 50 | Salience |
| Component 39 | 49% L superior frontal<br>16% L frontal orbital, 10% frontal operculum, 6% temporal pole, 6% inferior temporal (triangularis)<br>38% R supplementary motor (juxtapositional lobule) 20% paracingulate, 8% anterior cingulate | -12, 11, 71<br>-48, 18, -8<br>2, 6, 48 | Salience |
| Component 3 | 14% L planum temporale, 10% post-central gyrus, 7% superior temporal gyrus<br>49% L planum temporale, 10% parietal operculum, 6% central operculum | -66, -22, 14<br>64, -20, 12 | Auditory |
| Component 34 | 77% L frontal pole, 10% inferior frontal (triangularis)<br>37% L inferior frontal (triangularis) 27% middle frontal, 19% frontal pole<br>46% L supramarginal posterior, 14% supramarginal anterior | -48, 41, 11<br>-50, 34, 16<br>-50, -44, 54 | Language |
| Component 21 | 18% L frontal orbital, 17% inferior frontal, 10% frontal operculum<br>48% L superior lateral occipital, 37% angular gyrus<br>71% L inferior frontal (opercularis) 18% precentral | -51, 23, -7<br>-52, -62, 24<br>-56, 12, 8 | Language/Lateral Frontoparietal |
| Component 33 | 27% R superior lateral occipital, 25% angular gyrus<br>63% L superior lateral occipital, 10% angular<br>70% R angular, 7% superior lateral occipital<br>76% R middle frontal | 57, -58, 41<br>-50, -64, 46<br>60, -54, 26<br>46, 22, 44 | Bilateral Frontoparietal |
| Component 35 | 30% R superior lateral occipital<br>80% R frontal pole<br>24% R angular, 20% supramarginal posterior, 12% superior parietal lobule | 39, -61, 59<br>48, 44, 14<br>40, -48, 40 | Right Lateral Frontoparietal |
| Component 32 | 57% R frontal pole<br>75% R superior lateral occipital<br>65% R superior lateral occipital, 22% angular gyrus | 33, 65, -1<br>44, -70, 40<br>50, -60, 46 | Right Lateral Frontoparietal |
| Component 38 | 19% L superior lateral occipital<br>75% L frontal pole<br>37% L superior lateral occipital, 35% angular gyrus | -39, -64, 59<br>-36, 60, -4<br>-50, -60, 36 | Left Lateral Frontoparietal |
| Component 25 | 57% R occipital pole<br>68% L occipital pole | 21, -103, -4<br>-28, -98, -6 | Lateral Visual |
| Component 31 | 35% R inferior lateral occipital, 17% superior lateral occipital, 12% occipital pole<br>70% R superior lateral occipital<br>80% L inferior lateral occipital | 39, -85, 5<br>34, -82, 24<br>-44, -82, -8 | Lateral Visual |
| Component 19 | 59% R intracalcarine cortex, 7% supracalcarine cortex<br>44% R lingual gyrus, 5% occipital fusiform<br>50% L intracalcarine cortex | 9, -73, 14<br>14, -68, -2<br>-8, -72, 8 | Medial Visual |
| Component 7 | 52% R Crus II, 48% Crus I<br>79% L Crus I, 8% Crus VI | 36, -76, -38<br>-22, -73, -31 | Cerebellar |

### 3.3. Is the social network index associated with resting-state networks?

#### 3.3.1. Spatial maps of RSN Connectivity

Multivariate main effects of SN embeddedness were observed in cerebellar (IC7, p = 0.04) and frontoparietal (IC32, p = 0.012) networks (Figure 1) although were not explained at the univariate level. SN contacts showed associations with the connectivity of anterior default mode (IC29, p = 0.03) and lateral frontoparietal (IC32, p = 0.002 & IC33, p = 0.010, Figure 1) networks, while main effects of SN diversity were observed in the auditory (IC3, p = 0.014) language/frontoparietal (IC21, p = 0.026) anterior default mode (IC29, p = 0.003) and two frontoparietal (IC32, p = 0.022 and IC33, p= 0.006) networks. The anterior DMN (IC29) was highlighted in the univariate findings, showing a positive association with SN contacts, localized to the right anterior-medial frontal pole (BA10, β’s: 0.81, 0.73, all q < 0.04, Figure 3A, Table S3). Additionally, a significant cluster located in the right anterior frontal pole (BA 9/10) was negatively associated with both SN contacts (β’s: -1.34 and -1.27, all q < 0.05, Figure 3A, Table S3) and SN diversity (β range = -0.5640 to -0.6044, all q < 0.04, Figure 3B, Table S3).

**Figure 1:**
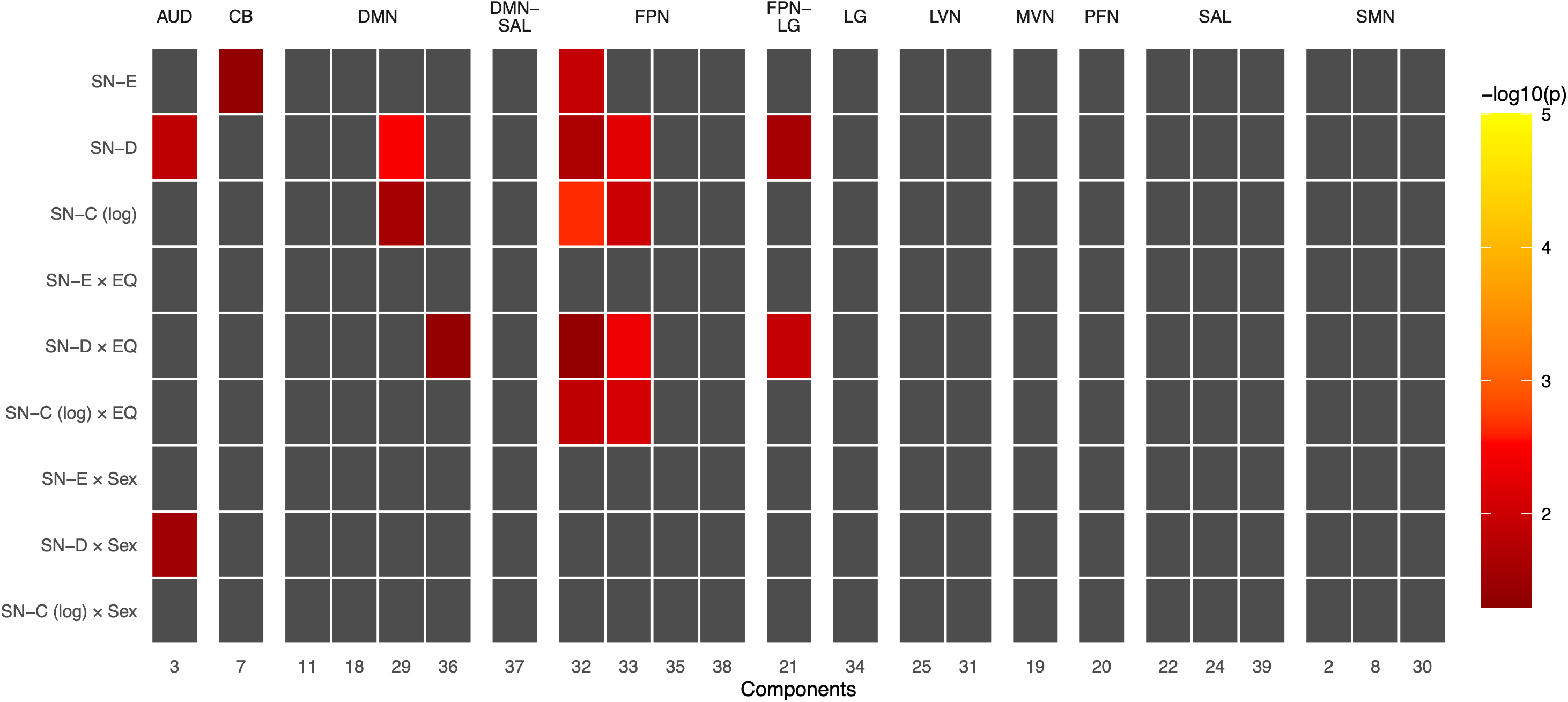
Multivariate associations between Social Network Index measures and their interactions with sex and Empathy Quotient across independent component connectivity spatial maps. *Legend:* Significant multivariate results (p < 0.05) across all three models, p value is log transformed. SN-E/-D/-C = social network embeddedness/diversity/contacts, EQ = empathy quotient. AUD = auditory network, CB = cerebellar network, DMN = default mode network, DMN-SAL = default mode-salience, FPN = frontoparietal network, LG = language, FPN-LG = frontoparietal-language, LVN = lateral visual network, MVN = medial visual network, SMN = sensorimotor network, PFN = pre-frontal network, SAL = salience network.

**Figure 2:**
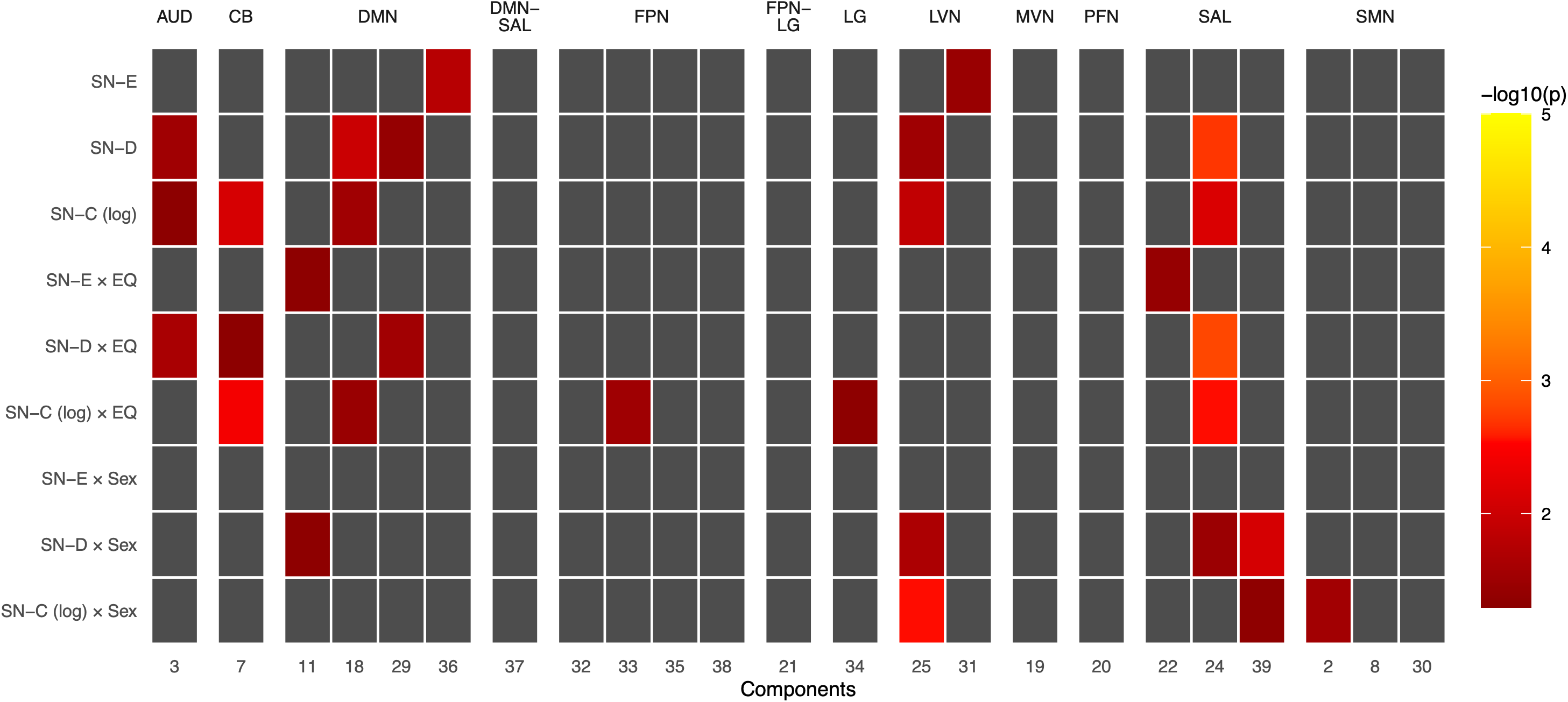
Multivariate associations between Social Network Index measures and their interactions with sex and Empathy Quotient across independent component power spectra. *Legend:* Significant multivariate results (p < 0.05) across all three models, p value is log transformed. SN-E/-D/-C = social network embeddedness/diversity/contacts, EQ = empathy quotient. AUD = auditory network, CB = cerebellar network, DMN = default mode network, DMN-SAL = default mode-salience, FPN = frontoparietal network, LG = language, FPN-LG = frontoparietal-language, LVN = lateral visual network, MVN = medial visual network, SMN = sensorimotor network, PFN = pre-frontal network, SAL = salience network.

**Figure 3.**
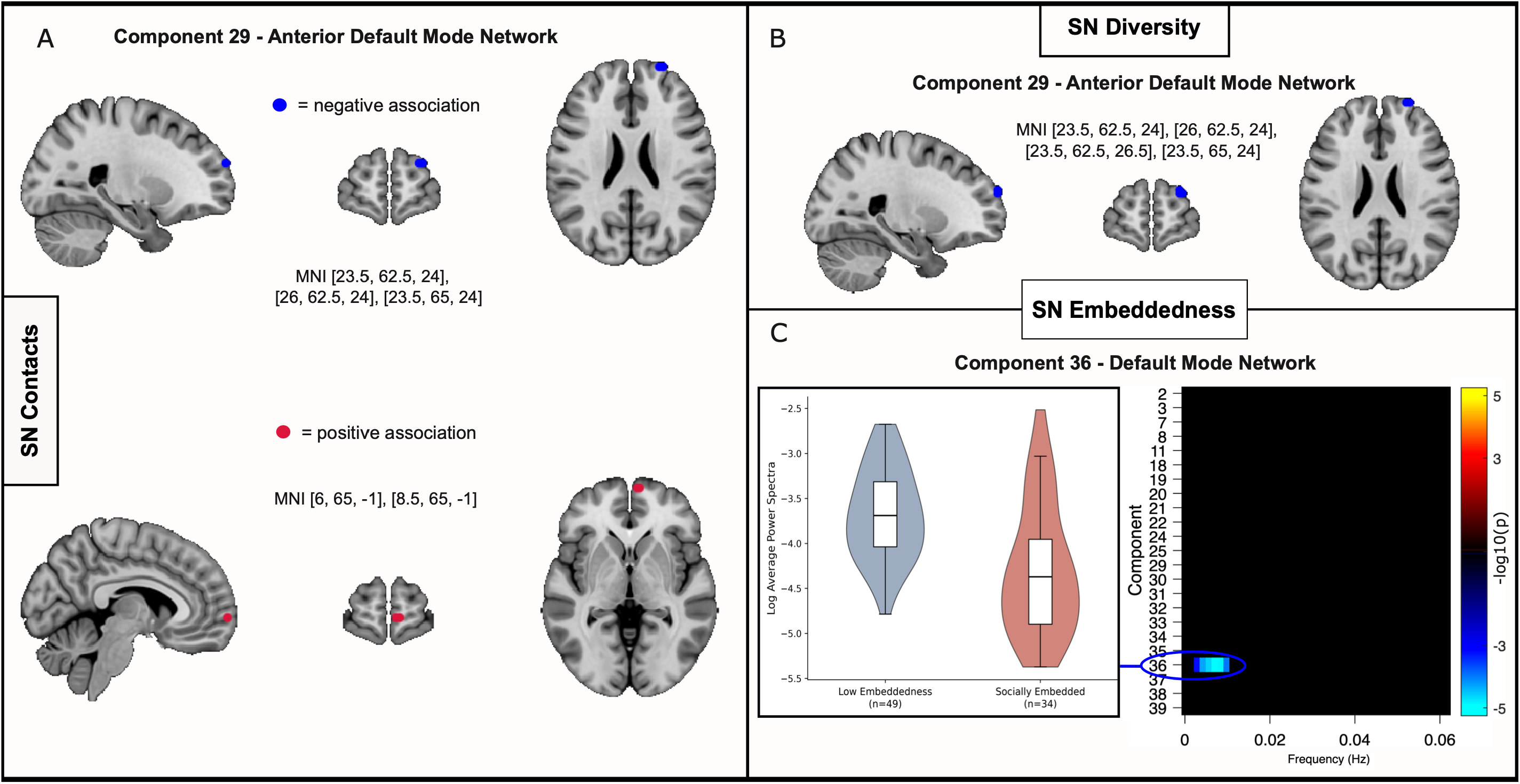
Default mode network connectivity and power associated with the Social Network Index. *Legend:* A) Spatial map associations with social network contacts, red indicates positive associations, blue indicates negative. B) Spatial map associations with social network diversity. C) Power spectra associations of the DMN with social network embeddedness, with the violin plot showing the average log(power) ± 1 SE for low embeddedness(blue) and socially embedded (red) p-values and average power are log-transformed. All findings shown survived leave-one-out sensitivity analysis (Table S4 and S5).

#### 3.3.2. RSN Power Spectra

Multivariate associations between both SN diversity and SN contacts with spectral power of the auditory (IC3; p = 0.03 and 0.047, respectively) default mode (IC18; p = 0.01 and 0.03) salience (IC24; p = 0.002 and 0.013) and lateral visual (IC25; p = 0.03 and 0.013) networks were observed (Figure 2). Additional multivariate effects showed SN diversity was uniquely associated with the anterior DMN (IC29, p = 0.04) whereas SN contacts were associated with the cerebellar network (IC7, p = 0.008, Figure 2). However, none of these associations were explained by univariate analyses.

In contrast, SN embeddedness was the only social metric to show univariate effects on power spectra. Multivariate associations were originally observed in lateral visual (IC31, p = 0.035) and default mode (IC36, p = 0.02) networks. Univariate findings implicated power of the DMN (IC36) as associated with SN embeddedness, where individuals classed as “Low Embeddedness” displayed higher power at very low frequencies (0.0028-0.0098 Hz) compared to socially embedded participants (β range = -0.420 to -0.677, Figure 3C).

### 3.4. Do the associations between social networks and resting-state networks differ according to empathizing ability?

#### 3.4.1. Spatial Maps of RSN Connectivity

Main multivariate effects of empathy on connectivity were observed in sensorimotor (IC2, p = 0.034) frontoparietal (IC20, p = 0.02) anterior default mode (IC29, p = 0.03) and default/salience (IC37, p = 0.02) networks indicating unique main effects regardless of SNI variables. Univariate analysis revealed a large cluster in the anterior default mode network (Table S3) located in the frontal pole, spanning the dorso-lateral pre-frontal cortex, superior frontal and middle frontal gyri (β range: 0.038-0.085, all q < 0.05) where increased connectivity was associated with higher empathy (Figure 4A) and an additional cluster in the inferior frontal pole/pars triangularis negatively associated with EQ (β = -0.022, q = 0.024; Table S2).

**Figure 4.**
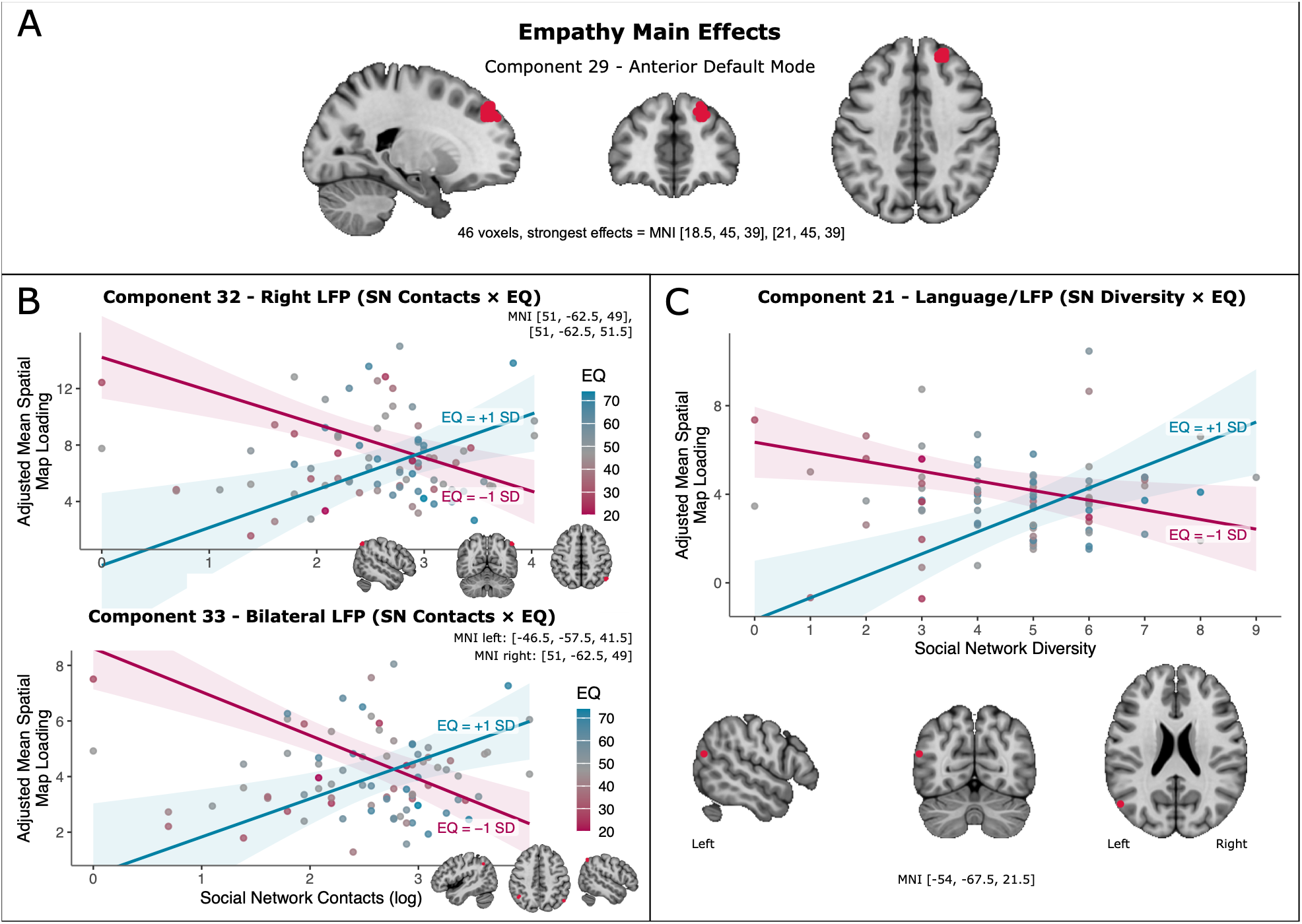
Functional Connectivity associated with Empathy and Social Network Indices. *Legend:* A) Main effects of empathy on connectivity of the anterior DMN. B) Significant interactions between empathy and social network (SN) contacts on spatial maps of connectivity on lateral frontoparietal components. Points represent each participant’s covariate-adjusted (sex, age, FWD, TR) mean z-score loadings across statistically significant voxels. Lines indicate model predicted loadings at low EQ (-1 SD) and high EQ (+1 SD) with covariates held at their mean. Shaded regions indicate 95% confidence intervals reflecting uncertainty in the estimated regression lines. Point color denotes continuous EQ score. C) Significant interactions between empathy and social network diversity on spatial maps of connectivty in the lateral frontoparietal/language component. All findings shown survived leave-one-out sensitivity analysis (Table S4).

Multivariate analysis of the interaction between empathy and SN contacts revealed effects in two lateral frontoparietal components (IC32, p = 0.0134 and IC33, p = 0.008). Interestingly, both components showed significant univariate effects in the same location (MNI: 51, -62.5, 49 and 51, -62.5, 51.5) spanning the right superior lateral occipital and angular gyri (IC32 β range: 0.178-0.261, IC33 β range: 0.174-0.179; Table S3). These findings consisted of individuals with higher EQ exhibiting a positive association between SN contacts and connectivity in these regions but a negative association in individuals with lower EQ (Figure 4B). In addition to this cluster, IC33 showed significant interactions in the left hemisphere, overlapping with angular, supramarginal and superior lateral occipital gyri (β range: 0.0473 to 0.0985, Figure 4B, Table S3).

Similarly to SN contacts, multivariate analyses detected a significant SN diversity and EQ interaction in the same two lateral frontoparietal (IC32, p = 0.039; IC33, p = 0.005) networks and additionally in the left language/frontoparietal (IC21, p = 0.012) and default mode (IC36, p = 0.04) networks. Univariate analyses identified a cluster in the language/frontoparietal network (IC21; Figure 4C, Table S3) where, consistent with previous results, the association between SN diversity and FC of the superior lateral occipital gyrus differed by empathy level (β = 0.053, all q < 0.03). Lower-empathy individuals showed higher superior lateral occipital connectivity associated with reduced social diversity, whereas individuals with increased empathy showed the opposite pattern with higher connectivity relating to more social diversity (Figure 4C). These interactions between SN indices and empathy on connectivity survived sensitivity analyses (Table S4).

#### 3.4.2. RSN Power Spectra

SN embeddedness showed significant interactions with empathy in the connectivity of posterior DMN (IC11, p = 0.047) and salience (IC22, p = 0.038) networks effects that were not explained by the univariate analyses. In contrast, relatively widespread effects were observed for SN contacts, with multivariate interactions between empathy and social contacts observed in power spectra of cerebellar (IC7, p = 0.0038) default mode (IC18, p = 0.035) lateral frontoparietal (IC33, p = 0.03) language (IC34, p = 0.046) and salience (IC24, p = 0.003) networks (Figure 2). Default mode (IC18) and salience (IC24) networks showed frequency-dependent SN contacts-by-EQ interactions (Figure 5A). Figure 5B shows the interaction within this range for the salience network; in particular within the range of 0.1088-0.1256 Hz the direction of the association between SN contacts and power differed by empathy level (Figure 5B). Among individuals with high empathy, greater social networks were associated with greater high-frequency power, while low empathy individuals exhibited a reversed pattern with greater social networks associated with lower high-frequency power (average beta = 0.026). This association was in the opposite direction at lower frequencies (0.0112-0.0209 Hz, average beta = -0.023). Finally, while the cerebellar network (IC07) was also significantly associated with the interaction between SN contacts and EQ, but this effect was not statistically robust (Table S5).

**Figure 5.**
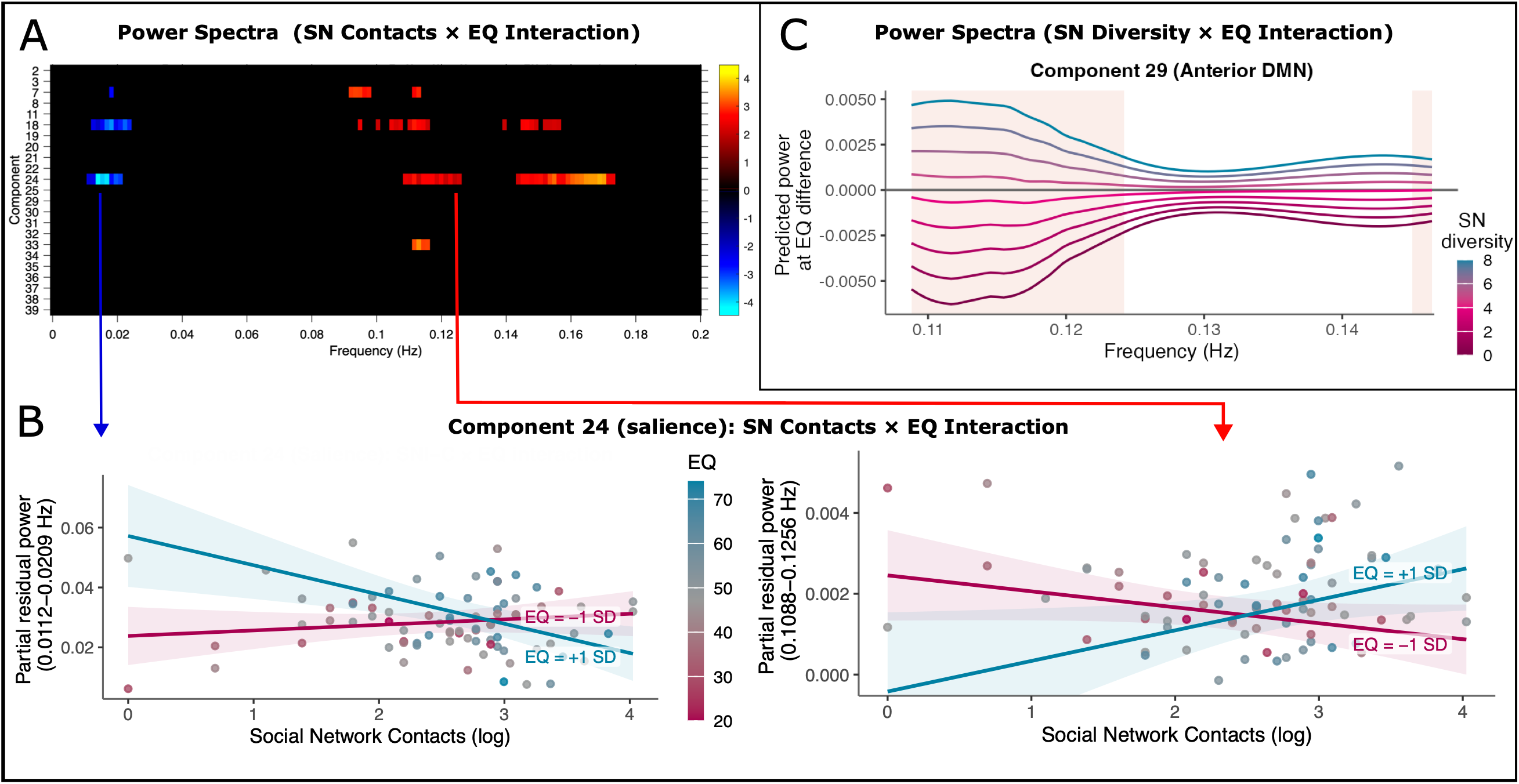
Power Spectra within the Default Mode and Salience networks associated with Empathy and Social Network Indices. *Legend:* A) Components where power bands demonstrated a significant interaction between social network (SN) contacts and empathy (EQ) color bar indicates logged FDR-corrected p-values B) Plots indicating frequency-specific power of the salience network against SN contacts separated by upper (61.6, blue) and lower (37.1, pink) standard deviations of the EQ (left most 0.0112-0.0209Hz, right most 0.1088-0.1256 Hz). Points represent residual power after adjusting for covariates (sex, age, TR, FWD). Y-axis represents average power across the frequency range with a data point per participant. Regression lines are based on individuals with one standard deviation above (61.6, blue) and one standard deviation below (37.1, pink) the mean EQ score. C) Each line represents a single social network diversity (SN diversity) score (0-8) and y-axis represents power at EQ difference (power at high EQ – power at low EQ). In individuals with greater social diversity, higher empathy was associated with greater power (blue lines). In individuals with lower social diversity, higher empathy was associated with lower power (maroon lines). The difference was largest at lower frequencies, with red shading indicating frequencies where significant effects were detected.

SN diversity and empathy interacted significantly in resting state network power of the auditory (IC3, p = 0.023) cerebellar (IC7, p = 0.048) salience (IC24, p = 0.0016) and anterior default mode (IC29, p = 0.029) networks. Univariate analyses showed SN diversity-by-EQ interactions at frequencies of 0.1102 to 0.1228 Hz (average beta = 0.0148) and 0.1451 Hz (beta = 0.0139) in the anterior DMN (IC29). In both frequency ranges, the association between empathy and power depended on SN diversity. In individuals with greater social diversity, higher empathy was associated with greater power (Figure 5C, blue lines). In individuals with lower social diversity, higher empathy was associated with lower power (Figure 5C, maroon lines). The difference was largest at lower frequencies.

### 3.5. Do the associations between social networks and resting-state networks differ between males and females?

#### 3.5.1. Spatial Maps of RSN Connectivity

Multivariate spatial map interactions were observed between SN diversity and sex in the auditory network only (IC3, p = 0.03; Figure 1). No spatial map interactions between any of the SNI metrics and sex survived at the univariate level.

#### 3.5.2. RSN Power Spectra

Multivariate power spectra findings implicated an SN contacts and sex interaction in sensorimotor (IC2, p = 0.028) lateral visual (IC25, p = 0.0028) salience (IC39, p = 0.045, Figure 2). Females showed a positive relationship between SN contacts and average power between 0.0670-0.1549 Hz of the salience network and differed significantly to that in males, suggesting salience network synchronicity relates to the number of social contacts in females more so than males (Figure 6A).

**Figure 6.**
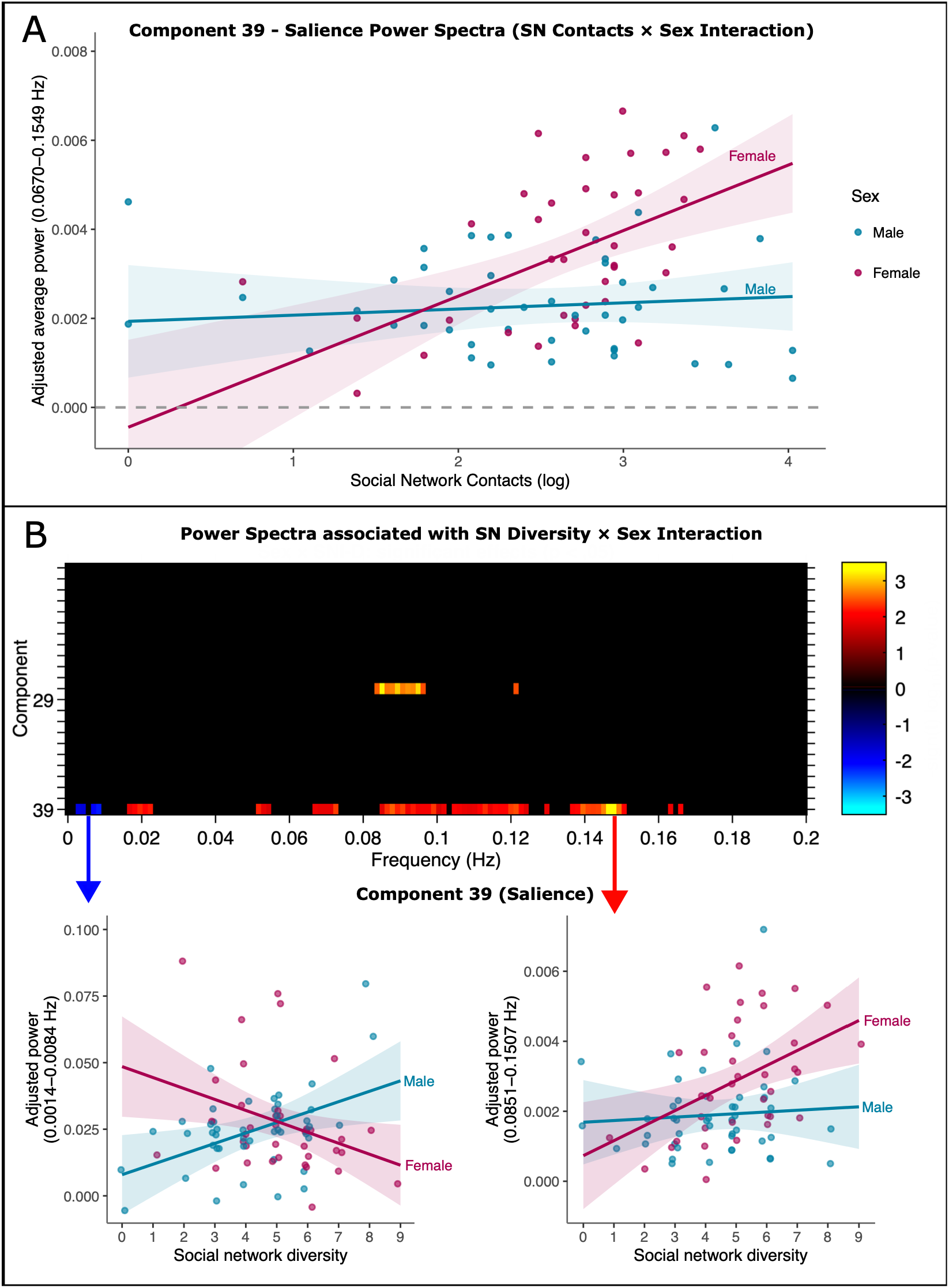
Power Spectra associated with Sex and Social Network Indices. *Legend:* A) Interaction between social network contacts (SN contacts) and sex on power spectra of the salience network (component 39). Points represent average residual power after adjusting for covariates (sex, age, TR, FWD) with a data point per participant. Lines show model-predicted power for each sex with 95% CIs. B) Top panel shows components with significant social network diversity-by-sex power spectra effects. Bottom panel shows the different interactions observed at each significant frequency band in component 39 (salience network). All findings shown survived leave-one-out sensitivity analysis (Table S5).

SN diversity interacted with sex at the multivariate level in posterior default mode (IC11, p = 0.046) lateral visual (IC25, p = 0.022) and salience (IC39, p = 0.008, IC24, p = 0.033) networks. Univariate analyses revealed associations with the lateral visual network (IC25) within 0.0837-0.0963 Hz range (β = 0.069) where a positive relationship between SN diversity and spectral power was present in females that differed significantly that in males. The salience network (IC39) additionally showed frequency-dependent associations, where at lower frequencies females showed a more negative association between SN diversity and power then males relative to high frequencies where the relationships were reversed and similar to those observed for SN contacts (Figure 6B). Finally, no interactions were observed between SN embeddedness and sex in both multivariate and univariate results.

## 4. Discussion

This study investigated how social network characteristics relate to resting-state brain connectivity and power in a healthy sample and determined the role of empathy and sex in these relationships. We demonstrate that social connection, indexed as the number of contacts an individual has and the variance in an individual’s social roles, is significantly related to connectivity of the default mode network, a network associated with internal mental processes. Moreover, greater variance in the social roles of those contacts (having different contact domains) termed social embeddedness, is associated with lower power at low frequencies of default mode network activation. Importantly, the neural associations with social connection differed depending upon a persons’ empathizing ability in the frontoparietal, salience and default mode networks. Finally, spectral power of the salience network, in particular, exhibited a differential relationship with social variables in males relative to females.

Power spectra capture the power distribution across frequencies variance in a components’ time course (Allen et al., 2011), however, the precise neurophysiological properties underlying power across resting-state spectra are poorly understood. Research suggests that a variety of both high and low frequencies contribute to the BOLD signal (Kajimura et al., 2023) and some theories propose that lower frequencies may reflect more long-range brain dynamics while higher frequencies may alternatively reflect overactive (Qiao et al., 2024) more locally specific brain activity (Pan et al., 2013). Spectral power therefore provides a novel perspective into the neural correlates of social networks; however, our understanding of its underlying mechanisms is still limited.

### 4.1. The role of the anterior default mode network in social networks

Across findings, it is evident that the anterior DMN represents a major correlate of both social network characteristics. Specifically, individuals with lower SN contacts and diversity showed greater dorsomedial prefrontal cortex (dmPFC) connectivity, and individuals with low social embeddedness showed higher low-frequency power in the DMN compared to socially embedded individuals, painting a cohesive picture of increased DMN connectivity relating to reduced social behaviors. These findings align with research linking the anterior DMN with social networks (Noonan et al., 2018) and broader literature showing greater DMN connectivity in individuals with self-reported loneliness (Spreng et al., 2020) and lower self-perceived social status (Muscatell et al., 2012). Structurally, the volume of the dmPFC has been negatively associated with social contacts (Veerareddy et al., 2023) and emotional intelligence (Takeuchi et al., 2011). Of note, one study found higher resting-state activation in the medial pre-frontal cortex of the anterior DMN associated with lower affective/emotional empathy (Oliveira Silva et al., 2018). Spreng *et al*. (2020) propose that reduced social interaction may contribute to a shift to internally directed social-cognitive processes, and the engagement of the dmPFC has been observed during tasks that emphasize self-awareness and self-reference during social interaction (Gusnard et al., 2001; Schilbach et al., 2006). Our findings may reflect a mechanism whereby individuals with low social contacts and diversity use internal reflection to compensate for the lack of external social stimuli (Cacioppo & Hawkley, 2009; Dunbar, 2025).

Conversely, the reduced connectivity observed in individuals with high social contacts may reflect more efficient social-cognitive processing (Takeuchi et al., 2011; Veerareddy et al., 2023). Notably, Veerareddy *et al*. (2023) previously found that the association between the volume of this region and social contacts was mediated by empathizing ability, and a *post-hoc* mediation analysis herein corroborates this, showing that the empathy quotient mediates the association between dmPFC connectivity and number of social contacts (Figure S2).

Nearby, anteromedial PFC (amPFC) demonstrates the opposite pattern, with higher connectivity associated with higher social contacts. The amPFC is a core node of the DMN known to play a role in self-referential processing, thinking about future and past events, and activates when making decisions related to individuals we care about and when understanding social meaning (Andrews-Hanna et al., 2010, 2014; Lieberman et al., 2019; Spreng & Andrews-Hanna, 2015). It is plausible that the positive association observed herein may reflect the ability to distinguish between the self, versus others, in social situations, as important for maintaining social contacts (Finlayson-Short et al., 2020; Oliveira Silva et al., 2018). These divergent findings across anatomically neighboring regions of the PFC suggest that social behaviors may differentially relate to coordination between cortically fine-tuned locations subserving socio-cognitive functions (W. Li et al., 2014).

### 4.2. Empathizing ability modulates the relationship between resting-state connectivity and social connection

In line with existing research (S. J. Kim et al., 2017; Oliveira Silva et al., 2018) we found an association between EQ and connectivity in the same anterior DMN component localized to the frontal pole, spanning the anterior dorsolateral PFC. Moreover, we observed that empathy additionally altered how SN diversity related to the spectral power and not spatial, characteristics of this network. This is consistent with Che *et al*. (2014)’s study showing higher DMN synchronicity in individuals with more social support, where the authors posit that DMN activity may be linked to social behaviors through empathizing and theory of mind abilities. The positive relationship observed in individuals with high-empathy may indicate more synchronization within higher frequency bands of the anterior DMN as a neurobiological process associated with empathy and social network diversity. For participants with low-empathy the direction of effect was reversed, where higher power related to low-empathy and low social diversity. It is worth noting that within similar frequency bands as our effect (0.1102-0.1228 Hz) higher power in individuals with schizophrenia (Garrity et al., 2007) and bipolar disorder (Dahan et al., 2026) has been observed in the DMN. Given that social (Blain et al., 2025; Isele et al., 1985) and empathetic (Derntl et al., 2009, 2012) deficits are frequently observed in these populations, future research examining how social behavior and empathy relate to default mode network power in psychiatric samples would address this potentially informative research gap.

Beyond the anterior DMN, the lateral frontoparietal network showed differing associations with our social measures depending on empathizing ability. Social network embeddedness did not interact with empathy, but both SN contacts and SN diversity exhibited distinct interactions with empathy on different components representing this network, suggesting these aspects of social behavior may depend more on relating to the feelings of others. These findings consistently implicated the right superior lateral-occipital cortex, a region which contributes to the perception of body motion (Murray, 2003) and human movement (Grosbras et al., 2012) and thickness of this region has been associated with cognitive empathy previously (Uribe et al., 2019). Interestingly, the empathy quotient has been linked to visual processing of body form (Miller & Saygin, 2013) and an MEG study found that individuals with high EQ scores showed greater right lateral occipital activation when observing social interactions, interpreted as an increased focus on facial and body cues (Hamada et al., 2022). Dzuira & Thompson (2014) also found that a nearby region of lateral occipitotemporal cortex was activated during perception of social body movements and that this activity correlated with social network diversity. Our findings compliment this work (Dziura & Thompson, 2014) and taken together suggest that functional connectivity at rest in this region may relate to social network maintenance via empathetic perception of non-verbal social cues The lateral frontoparietal network clusters associated with SN contacts also included bilateral angular gyri, a core region associated with theory of mind (Mar, 2011) and the adjacent area of the temporoparietal junction is crucial for distinction of the self and others during empathetic response (Lamm et al., 2019). The overlap of these spatial map clusters with regions crucial for theory of mind and higher-order visual processing may reflect the integrated contribution of social visual processing (such as body language) and theory of mind in maintaining social networks, where visual social cues can help to infer mental states of others.

Additionally, empathy significantly moderated how SN contacts related to the spectral power of the DMN, salience, and bilateral frontoparietal networks, three networks that have previously been linked to emotional empathy (Bilevicius et al., 2018). Frequency band specific effects were observed in the DMN (IC18) and salience (IC24) networks; at higher frequencies SN contacts was negatively associated with power in individuals with low-empathy but positively associated in those with high-empathy. Here, increased high frequency power was observed in both high-empathy individuals with more social contacts and low-empathy individuals with less social contacts, showing that a one-size-fits-all understanding is not applicable in this context.

Although difficult to interpret, this pattern of a positive relationship high-empathy individuals but the opposite in individuals with low-empathy may be consistent with the differential susceptibility theory (Ellis et al., 2011). This theory states that individuals vary in how their social and physical environments affect them, such that factors that may be positive for some individuals (such as more social contacts) may not always produce positive outcomes (Assary et al., 2023). In fact, variability in environmental sensitivity has been proposed to partly derive from brain networks supporting attention, emotional response and empathy (Assary et al., 2023) such as the salience (which overlaps with regions implicated in empathy ) default, and lateral frontoparietal networks (Homberg & Jagiellowicz, 2022) consistent with those implicated herein. Understood in this framework, the opposing relationships observed at different levels of empathy may relate to differential environmental sensitivity across individuals. For instance, similar to the relationship between social contacts and depression risk described by Dunbar (2025) the same number of contacts might be cognitively resourceful for high-empathy individuals but cognitively demanding for those with lower empathy. Alternatively, although higher empathy is generally believed to be beneficial, increased empathetic tendencies can also lead to maladaptive outcomes (Tone & Tully, 2014) and our findings could alternatively reflect high empathy and increased social networks being more cognitively demanding.

Evidently, how resting-state connectivity relates to social networks is shaped by one’s ability to relate to others however, the present study is limited by its cross-sectional nature, and the causal direction of all effects cannot be inferred. Nevertheless, this research provides an avenue for future investigations in both psychiatry and social health, where research and interventions targeting individual differences in empathizing ability and psychological disposition may be helpful in understanding the neurobiology of social deficits and developing social health interventions (Rauchbauer et al., 2023).

### 4.3. The influence of birth sex on social network and brain function associations

We observed that sex moderated how social networks relate to the power of resting-state networks at certain frequency bands, however effects were observed in their patterns of spatial connectivity. While no studies have investigated these relationships previously, SN-contacts and SN-diversity interacted with sex in their association with the power of the salience and lateral visual networks (Figure 6). In both cases, at higher frequencies greater social scores were associated with higher power in females relative to males, potentially suggesting that higher frequency power of intrinsic brain networks is particularly relevant for social contacts and diversity in females. The salience network plays a role in diverting attention towards salient stimuli (Uddin, 2016; Uddin et al., 2019), while the lateral visual network is involved in visual processing. Potentially consistent with this finding, existing studies have found sex differences in salience network connectivity (Allen et al., 2011; S. Zhang et al., 2025) and higher connectivity in both of these networks has been previously associated with social connection (Pillemer et al., 2017; Pruitt et al., 2024).

Variability in social network characteristics across sexes is well-documented (David-Barrett et al., 2015; Dunbar et al., 2024; Williams et al., 2022) and SN properties may differently relate to well-being across sexes, for example, Shin and Park (2023) found that group diversity in their sample was more critical for well-being in females. Therefore, the same level of social diversity and contacts may provide different cognitive and emotional demands across sexes (Farago et al., 2026). The implication of networks involved in attention and visual processing herein may be relevant to studies showing that females tend to exhibit stronger visual social-attention abilities (L. Zhang et al., 2025) and emotion recognition (Christov-Moore et al., 2014) as well as more efficient processing of, and greater interest in, social stimuli (Proverbio, 2017, 2023; Proverbio et al., 2008).

It is plausible that empathy and sex may also jointly moderate the association between social relationships with connectivity, especially given females tend to report higher empathy levels than males (Christov-Moore et al., 2014). However, the current sample was too small for the investigation of three-way interactions (e.g. there are only 5 females with low empathy and high social embeddedness). Additionally, causal inferences were not possible in this study; it is equally likely that sex-related differences in brain function may shape social behavior, and/or environmental differences related to sex socialization may shape both social behavior and brain connectivity patterns. Nonetheless, these findings provide further support for sex differences in the relationships between social behavior and brain function, in particular, variability in network power.

### 4.4. Social network embeddedness and resting-state networks

As discussed above, the only univariate finding implicating SN embeddedness indicated higher power in the DMN in individuals with limited social embeddedness. While SN contacts and diversity capture network characteristics more related to oneself (how many contacts and social roles one has) and were highly correlated with each other (Figure S1) social embeddedness captures properties related to the individuals’ contacts’ roles rather than their own and was less statistically related to ‘contacts’ or ‘diversity’ (Figure S1). This may suggest that embeddedness captures a different feature of a persons’ social network. However, whether the more indirect nature of this measure explains the reduced brain associations, remains unclear. Although the SN embeddedness sub-measure has been associated with amygdala volume and connectivity in previous studies (Bickart et al., 2011, 2012) many existing investigations linking the SNI to imaging-derived-phenotypes do not use this measure in their study (Blumen & Verghese, 2019; Molesworth et al., 2015; Pillemer et al., 2017) leaving limited prior evidence against which to interpret these findings.

## 5. Conclusions

To our knowledge, this is the first study to examine whether empathy or sex moderate associations between different social network characteristics and resting-state brain function, and to consider these associations using both spectral power and connectivity of resting-state networks. Overall, the accumulated findings from this study allow for three core conclusions. Social contacts and diversity shared many common correlates, including anterior default mode connectivity and strong interactions with empathy and sex, while social network embeddedness was independently associated the spectral power of the DMN. Secondly, the relationship between multiple social network characteristics and connectivity in the superior lateral occipital/temporo-parietal junction regions differed by empathizing ability, revealing a complex interplay between empathy, real-world social networks, and theory-of-mind/visual cue perception specialized regions. Finally, social network characteristics differentially related to network oscillations in lateral visual and salience networks in males and females, corroborating findings of unique neural underpinnings of social behavior between birth sexes. These observations highlight the importance of considering individual differences in empathy as well as sex-effects when studying brain and social behavior relationships, with implications for understanding individual differences in social tendencies and potential benefits if applied to better understand social deficits in psychiatric populations.

## Acknowledgements

The authors would like to thank the team at the California Institute of Technology for collecting this dataset and making it openly available on OpenNeuro (Kliemann et al., 2022) and the participants who made this work possible.

## Declarations

### Funding

This research was funded by Taighde Eireann (Research Ireland) proposal number 21/FFP-P/10165.

### Conflicts of Interest

No conflicts of interest to declare.

### Author Contributions

Author contributions included conceptualization and study design (MGC, AF, DMC) funding acquisition (DWM, DMC) supervision (DMC) data curation (MGC) methodology (MGC, AF) formal analysis (MGC, AS) analysis support (AF, DMC) results interpretation (AF, LR, AS, TV, AL, DMC) writing the original draft, visualization and administration (MGC) revising and editing (MGC, AL, DMC) approval and edits of the final version for publication (all authors).

## Notes

### Competing Interest Statement

The authors have declared no competing interest.

