## Supplementary_Data for "Social Connection: Resting-State Functional Connectivity Associations with Social Networks, and the Roles of Empathy and Sex"

### Supplementary Methods

**fMRIPrep Processing:** The data used in this study were extracted from OpenNeuro (ds003798) as minimally preprocessed derivatives. The description of the fMRIPrep processing used by Kliemann et al. (2022) is adapted below. FMRIprep generates a range of outputs and confounds; for the purpose of this study only the volumetric resting-state BOLD data in MNI152NLin2009cAsym space were used. Of the confounds generated, only framewise displacement was used as a covariate in the models.

**Supplementary Box 1.** Preprocessing description generated by fMRIPrep 20.2.1

Results included in this manuscript come from preprocessing performed using fMRIPrep 20.2.1 (Esteban et al., 2019; Esteban et al., 2018; RRID:SCR_016216), which is based on Nipype 1.5.1 (Gorgolewski et al., 2011; Gorgolewski et al., 2018; RRID:SCR_002502).

***Anatomical data preprocessing***

A total of 1 T1-weighted (T1w) images were found within the input BIDS dataset. The T1-weighted (T1w) image was corrected for intensity non-uniformity (INU) with `N4BiasFieldCorrection` (Tustison et al., 2010), distributed with ANTs 2.3.3 (Avants et al., 2008; RRID:SCR_004757), and used as T1w-reference throughout the workflow. The T1w-reference was then skull-stripped with a *Nipype* implementation of the `antsBrainExtraction.sh` workflow (from ANTs), using OASIS30ANTs as target template. Brain tissue segmentation of cerebrospinal fluid (CSF), white-matter (WM) and gray-matter (GM) was performed on the brain-extracted T1w using `fast` (FSL 5.0.9, RRID:SCR_002823; Zhang et al., 2001). Brain surfaces were reconstructed using `recon-all` (FreeSurfer 6.0.1, RRID:SCR_001847; Dale et al., 1999), and the brain mask estimated previously was refined with a custom variation of the method to reconcile ANTs-derived and FreeSurfer-derived segmentations of the cortical gray-matter of Mindboggle (RRID:SCR_002438; Klein et al., 2017). Volume-based spatial normalization to two standard spaces (MNI152NLin2009cAsym, MNI152NLin6Asym) was performed through nonlinear registration with `antsRegistration` (ANTs 2.3.3), using brain-extracted versions of both T1w reference and the T1w template. The following templates were selected for spatial normalization: *ICBM 152 Nonlinear Asymmetrical template version 2009c* (Fonov et al., 2009; RRID:SCR_008796; TemplateFlow ID: MNI152NLin2009cAsym), *FSL's MNI ICBM 152 non-linear 6th Generation Asymmetric Average Brain Stereotaxic Registration Model* (Evans et al., 2012; RRID:SCR_002823; TemplateFlow ID: MNI152NLin6Asym).

***Functional data preprocessing***

For each of the 4 BOLD runs found per subject (across all tasks and sessions), the following preprocessing was performed. First, a reference volume and its skull-stripped version were generated by aligning and averaging 1 single-band references (SBRefs). A B0-nonuniformity map (or *fieldmap*) was estimated based on two (or more) echo-planar imaging (EPI) references with opposing phase-encoding directions, with `3dQwarp` (Cox & Hyde, 1997; AFNI 20160207). Based on the estimated susceptibility distortion, a corrected EPI (echo-planar imaging) reference was calculated for a more accurate co-registration with the anatomical reference. The BOLD reference was then co-registered to the T1w reference using `bbregister` (FreeSurfer) which implements boundary-based registration (Greve & Fischl, 2009). Co-registration was configured with six degrees of freedom. Head-motion parameters with respect to the BOLD reference (transformation matrices, and six corresponding rotation and translation parameters) are estimated before any spatiotemporal filtering using `mcflirt` (FSL 5.0.9; Jenkinson et al., 2002). First, a reference volume and its skull-stripped version were generated using a custom methodology of *fMRIPrep*. The BOLD time-series were resampled onto the following surfaces (FreeSurfer reconstruction nomenclature): *fsaverage6*, *fsaverage*. The BOLD time-series (including slice-timing correction when applied) were resampled onto their original, native space by applying a single, composite transform to correct for head-motion and susceptibility distortions. These resampled BOLD time-series will be referred to as *preprocessed BOLD in original space*, or just *preprocessed BOLD*. The BOLD time-series were resampled into standard space, generating a *preprocessed BOLD run in MNI152NLin2009cAsym space*. First, a reference volume and its skull-stripped version were generated using a custom methodology of *fMRIPrep*. *Grayordinates* files (Glasser et al., 2013) containing 91k samples were also generated using the highest-resolution `fsaverage` as intermediate standardized surface space. Several confounding time-series were calculated based on the *preprocessed BOLD*: framewise displacement (FD), DVARS and three region-wise global signals. FD was computed using two formulations following Power (absolute sum of relative motions, Power et al., 2014) and Jenkinson (relative root mean square displacement between affines, Jenkinson et al., 2002). FD and DVARS are calculated for each functional run, both using their implementations in *Nipype* (following the definitions by Power et al., 2014). The three global signals are extracted within the CSF, the WM, and the whole-brain masks. The head-motion estimates calculated in the correction step were also placed within the corresponding confounds file. The confound time series derived from head motion estimates and global signals were expanded with the inclusion of temporal derivatives and quadratic terms for each (Satterthwaite et al., 2013). Frames that exceeded a threshold of 0.5 mm FD or 1.5 standardised DVARS were annotated as motion outliers. All resamplings can be performed with *a single interpolation step* by composing all the pertinent transformations (i.e. head-motion transform matrices, susceptibility distortion correction when available, and co-registrations to anatomical and output spaces). Gridded (volumetric) resamplings were performed using `antsApplyTransforms` (ANTs), configured with Lanczos interpolation to minimize the smoothing effects of other kernels (Lanczos, 1964). Non-gridded (surface) resamplings were performed using `mri_vol2surf` (FreeSurfer).

Many internal operations of *fMRIPrep* use *Nilearn* 0.6.2 (Abraham et al., 2014; RRID:SCR_001362), mostly within the functional processing workflow. For more details of the pipeline, see the section corresponding to workflows in *fMRIPrep*'s documentation (https://fmriprep.readthedocs.io/en/latest/workflows.html).

### Supplementary Results

**Table S1.** Pearson correlations between Social Network Indices and the Empathy Quotient.

| Variable | SN-Embed | SN-Contacts | SN-Diversity |
|---|---|---|---|
| **SN-Embed** | - | | |
| **SN-Contacts** | 0.68, $p = 1.946 \times 10^{-12}$ | - | |
| **SN-Diversity** | 0.64, $p = 7.76 \times 10^{-11}$ | 0.86, $p = < 2.2 \times 10^{-16}$ | - |
| **EQ** | 0.19, p = 0.08 | 0.28, p = 0.009 | 0.28, p = 0.009 |

*Legend:* Pearson's *r* correlation coefficients for the relationships between the SNI and the empathy quotient (n = 83). SN-Contacts was log-transformed. Correlations involving SN-Embed are point-biserial with SN-Embed coded 0 = limited embeddedness, 1 = socially embedded. SN = social network, EQ = empathy quotient.

**Table S2.** Participants demographic variables relationship with Social Network Indices and Empathy Quotient.

| | SNI-E (% Low, n Low) | Mean SNI-D (±SD) | Mean SNI-C (±SD) | Mean EQ(±SD) |
|---|---|---|---|---|
| **Total Sample** | 59.04%, (n = 49) | 4.64 (+/- 1.79) | 15.89 (+/- 14.21) | 49.37 (+/-12.23) |
| **Age (mean +/- SD)** | W = 715.5, p = 0.28 | ρ = 0.05, p = 0.64 | ρ = 0.03, p = 0.77 | ρ = 0.20, p = 0.07 |
| 29.29 +/-6.96 | | | | |
| **Sex at Birth** | $\chi^2$= 0.36, p = 0.55 | **W = 627.5, p = 0.04*** | W = 727, p = 0.26 | **W = 550, p = 0.006*** |
| Female (n = 37, 44.58%) | 54.05% (n = 20) | 5.11 (+/- 1.65) | 15.27 (+/- 7.33) | 53.30 (+/- 11.23) |
| Male (n = 46, 55.42%) | 63.04% (n = 29) | 4.26 (+/- 1.82) | 16.39 (+/- 18) | 46.21 (+/- 12.21) |
| **Race** | Fisher exact test, p = 0.52 | H = 2.66, p = 0.62 | H = 1.47, p = 0.83 | H = 4.04, p = 0.40 |
| African American/Black (n = 8) | 75% (n = 6) | 4.0 (+/- 1.41) | 15.25 (+/- 11) | 47.75 (+/- 13) |
| Asian (n = 12) | 41.67% (n = 5) | 5.0 (+/- 2.13) | 22.42 (+/-27.19) | 46.5 (+/- 10.14) |
| Caucasian/White (n = 37) | 56.76% (n = 21) | 4.81 (+/- 1.79) | 14.11 (+/-8.32) | 52.16 (+/- 12.00) |
| Multiracial (n = 17) | 70.59% (n =12) | 4.35 (+/- 1.62) | 14.59 (+/- 12.33) | 48.8 (+/- 13.46) |
| Other (n = 9) | 55.56% (n = 5) | 4.56 (+/- 2.01) | 17.56 (+/- 15.65) | 44.22 (+/- 12.21) |
| **Highest Education Level** | Fisher exact test, p = 0.61 | H = 1.68, p = 0.80 | H = 5.98, p = 0.201 | H = 2.57, p = 0.63 |
| High School (n = 4) | 75% (n = 3) | 3.75 (+/- 1.71) | 9.5 (+/- 6.03) | 44.75 (+/- 19.00) |
| Some College (n = 25) | 68% (n = 17) | 4.44 (+/- 2.14) | 15.08 (+/- 20.46) | 48.80 (+/- 12.32) |
| Associate's degree (n = 10) | 50% (n = 5) | 4.70 (+/- 1.49) | 17.3 (+/- 11.86) | 53.00 (+/- 15.37) |
| Bachelor's Degree (n = 37) | 51.35% (n = 19) | 4.78 (+/- 1.67) | 17.51 (+/-10.94) | 48.50 (+/- 11.20) |
| Postgraduate Degree (n = 7) | 71.43% (n = 5) | 5.00 (+/- 1.63) | 11.86 (+/-8.19) | 53.71 (+/- 9.41) |

*Legend:* Table shows the breakdown of each core demographic by each Social Network Index (SNI) measure and the Empathy Quotient (EQ). Statistics in shaded rows indicate whether a significant association is observed between the demographic variable and each IV (i.e.. is age associated with SNI-D?). SNI-E is a binary variable therefore the % and number of individuals classified as "Low embeddedness" is shown (i.e.. 20 of 37 females are classed as low social embeddedness). For continuous variables, the mean score per demographic is shown (i.e.. Females have a mean EQ of 53.30). Bold and * indicates a significant effect, here there was a significant difference in SNI-D and empathy across males and females

**Table S3.** Social Network Indices and their interactions with Empathy on independent component connectivity: spatial map location of significant effects in MNI space.

| Component | Direction of Effect | MNI co-ordinates (x, y, z) | Significance (q-value) |
|---|---|---|---|
| **Main effects of Social Network Diversity** | | | |
| Anterior Default Mode (IC29) | Negative | 23.5, 62.5, 24 | 0.023 |
| | | 26, 62.5, 24 | 0.027 |
| | | 23.5, 62.5, 26.5 | 0.020 |
| | | 23.5, 65, 24 | 0.031 |
| **Main effects of Social Network Contacts** | | | |
| Anterior Default Mode (IC29) | Positive | 6, 65, -1 | 0.031 |
| | | 8.5, 65, -1 | 0.011 |
| | Negative | 23.5, 62.5, 24 | 0.011 |
| | | 26, 62.5, 24 | 0.031 |
| | | 23.5, 65, 24 | 0.042 |
| **Main effects of the Empathy Quotient** | | | |
| Anterior Default Mode (IC29) | Positive | 13.5, 52.5, 19 | 0.024 |
| | | 13.5, 55, 19 | 0.032 |
| | | 21, 42.5 , 34 | 0.024 |
| | | 23.5, 42.5, 34 | 0.030 |
| | | 21, 45, 34 | 0.024 |
| | | 23.5, 45, 34 | 0.024 |
| | | 23.5, 47.5, 34 | 0.046 |
| | | 23.5, 50, 34 | 0.032 |
| | | 21, 52.5 , 34 | 0.030 |
| | | 23.5, 52.5, 34 | 0.026 |
| | | 18.5, 42.5, 36.5 | 0.040 |
| | | 21, 42.5 , 36.5 | 0.024 |
| | | 23.5, 42.5, 36.5 | 0.032 |
| | | 21, 45, 36.5 | 0.014 |
| | | 23.5, 45 , 36.5 | 0.014 |
| | | 21, 47.5 , 36.5 | 0.024 |
| | | 23.5, 47.5, 36.5 | 0.024 |
| | | 23.5, 50 , 36.5 | 0.038 |
| | | 18.5, 42.5, 39 | 0.046 |
| | | 21, 42.5 , 39 | 0.024 |
| | | 21, 45, 39 | 0.014 |
| | | 23.5, 45 , 39 | 0.024 |
| | | 21, 47.5 , 39 | 0.024 |
| | | 23.5, 47.5, 39 | 0.030 |
| | | 21, 45, 41.5 | 0.024 |
| | | 23.5, 45, 41.5 | 0.024 |
| | | 21, 47.5 , 41.5 | 0.024 |
| | | 23.5, 42.5, 44 | 0.029 |
| | | 21, 45, 44 | 0.024 |
| | | 23.5, 45 , 44 | 0.024 |
| | | 18.5, 32.5, 46.5 | 0.040 |
| | | 18.5, 35 , 46.5 | 0.046 |
| | | 23.5, 42.5, 46.5 | 0.024 |
| | | 21, 45, 46.5 | 0.024 |
| | | 23.5, 45 , 46.5 | 0.029 |
| | | 21, 37.5 , 49 | 0.030 |

| | | | |
|---|---|---|---|
| | | 21, 40, 49 | 0.029 |
| | | 23.5, 40 , 49 | 0.025 |
| | | 21, 42.5 , 49 | 0.024 |
| | | 23.5, 42.5, 49 | 0.024 |
| | | 21, 45, 49 | 0.026 |
| | | 21, 40, 51.5 | 0.024 |
| | | 21, 42.5 , 51.5 | 0.032 |
| | Negative | 46, 40, 11.5 | 0.024 |
| | | 46, 42.5, 11.5 | 0.048 |
| **Effects of the Interaction between Social Network Diversity and Empathy** | | | |
| L Language/Lateral Frontoparietal (IC21) | Positive | -54, -67.5, 21.5 | 0.023 |
| | | -51.5, -67.5, 21.5 | 0.023 |
| **Effects of the Interaction between Social Network Contacts and Empathy** | | | |
| Lateral Frontoparietal (IC32) | Positive | 51, -65, 49 | 0.019 |
| | | 51, -62.5, 49 | 0.009 |
| | | 53.5, -62.5, 49 | 0.024 |
| | | 48.5, -65, 51.5 | 0.028 |
| | | 51, -62.5, 51.5 | 0.024 |
| | | 48.5, -62.5, 49 | 0.034 |
| Lateral Frontoparietal (IC33) | Positive | -46.5, -57.5, 41.5 | 0.035 |
| | | -44, -55, 41.5 | 0.044 |
| | | -46.5, -60, 41.5 | 0.044 |
| | Positive | 51, -62.5, 49 | 0.035 |
| | | 51, -62.5, 51.5 | 0.035 |

*Legend:* Significant effects of the SNI measures and their interactions with empathy on individual voxels within resting-state network spatial maps. Q-values indicate FDR corrected p-values. All of these effects survived sensitivity analysis (see Table S3).

**Table S4.** Leave-one-out sensitivity analysis results for effects of the Social Network Index and Empathy Quotient on Spatial Maps.

| Phenotype | Component | Number of Voxels | Effect Direction | Minimum Coefficient | Maximum Coefficient | Maximum p-value | Effect survival |
|---|---|---|---|---|---|---|---|
| SNI-D | IC29 | 4 | Negative | -0.644 | -0.530 | $7 \times 10^{-6}$ | True |
| SNI-C | IC29 | 3 | Negative | -1.543 | -1.124 | $2.5 \times 10^{-5}$ | True |
| SNI-C | IC29 | 2 | Positive | 0.730 | 0.821 | $1.3 \times 10^{-5}$ | True |
| EQ | IC29 | 43 | Positive | 0.057 | 0.061 | $< 1.0 \times 10^{-6}$ | True |
| EQ | IC29 | 2 | Negative | -0.026 | -0.021 | $1.5 \times 10^{-4}$ | True |
| SNI-C × EQ | IC32 | 6 | Positive | 0.172 | 0.240 | $7.2 \times 10^{-4}$ | True |
| SNI-C × EQ | IC33 | 5 | Positive | 0.067 | 0.131 | 0.002 | True |
| SNI-D × EQ | IC21 | 2 | Positive | 0.040 | 0.051 | $1.2 \times 10^{-4}$ | True |

*Legend:* For each significant phenotype-component effect a leave-one-out analysis was conducted where models were refit removing a different participant each time, focusing on any participants with high Cook's distance. Effects were considered to survive where the maximum p-value remained significant when removing high influence participants.

**Table S5.** Leave-one-out sensitivity analysis results for the effects of the Social Network Index, Sex, and Empathy Quotient on Power.

| Phenotype | Component | Number of Frequencies | Effect Direction | Minimum Coefficient | Maximum Coefficient | Maximum p-value | Effect survival |
|---|---|---|---|---|---|---|---|
| SNI-E | IC36 | 7 | Negative | -0.691 | -0.615 | $1.30 \times 10^{-5}$ | True |
| SNI-D × EQ | IC29 | 10 | Positive | 0.012 | 0.016 | 0.004 | True |
| SNI-C × EQ † | IC07 | 8 | Positive | 0.020 | 0.038 | 0.067† | False |
| SNI-C × EQ | IC07 | 1 | Negative | -0.029 | -0.018 | 0.034 | True |
| SNI-C × EQ | IC24 | 36 | Positive | 0.028 | 0.035 | 0.004 | True |
| SNI-C × EQ | IC24 | 8 | Negative | -0.026 | -0.014 | 0.026 | True |
| SNI-C × EQ | IC18 | 20 | Positive | 0.020 | 0.030 | 0.024 | True |
| SNI-C × EQ | IC18 | 9 | Negative | -0.027 | -0.016 | 0.013 | True |
| SNI-C × EQ | IC33 | 5 | Positive | 0.027 | 0.034 | 0.011 | True |
| SNI-D × Sex | IC25 | 12 | Positive | 0.253 | 0.336 | 0.005 | True |
| SNI-D × Sex | IC39 | 55 | Positive | 0.193 | 0.278 | 0.002 | True |
| SNI-D × Sex | IC39 | 5 | Negative | -0.280 | -0.165 | 0.042 | True |
| SNI-C × Sex | IC02 | 10 | Positive | 0.632 | 0.885 | 0.003 | True |
| SNI-C × Sex | IC39 | 75 | Positive | 0.531 | 0.852 | 0.004 | True |

*Legend:* For each significant phenotype-component effect a leave-one-out analysis was conducted where models were refit removing a different participant each time, focusing particularly on any participants with high Cook's distance. Effects were considered to survive where the maximum p-value remained significant when removing high influence participants. † = indicates effects that did not survive and were likely driven by high influence participants.

**Figure S2.** Empathy Quotient mediated the association between social network contacts and mean connectivity of the right dorsomedial pre-frontal cortex

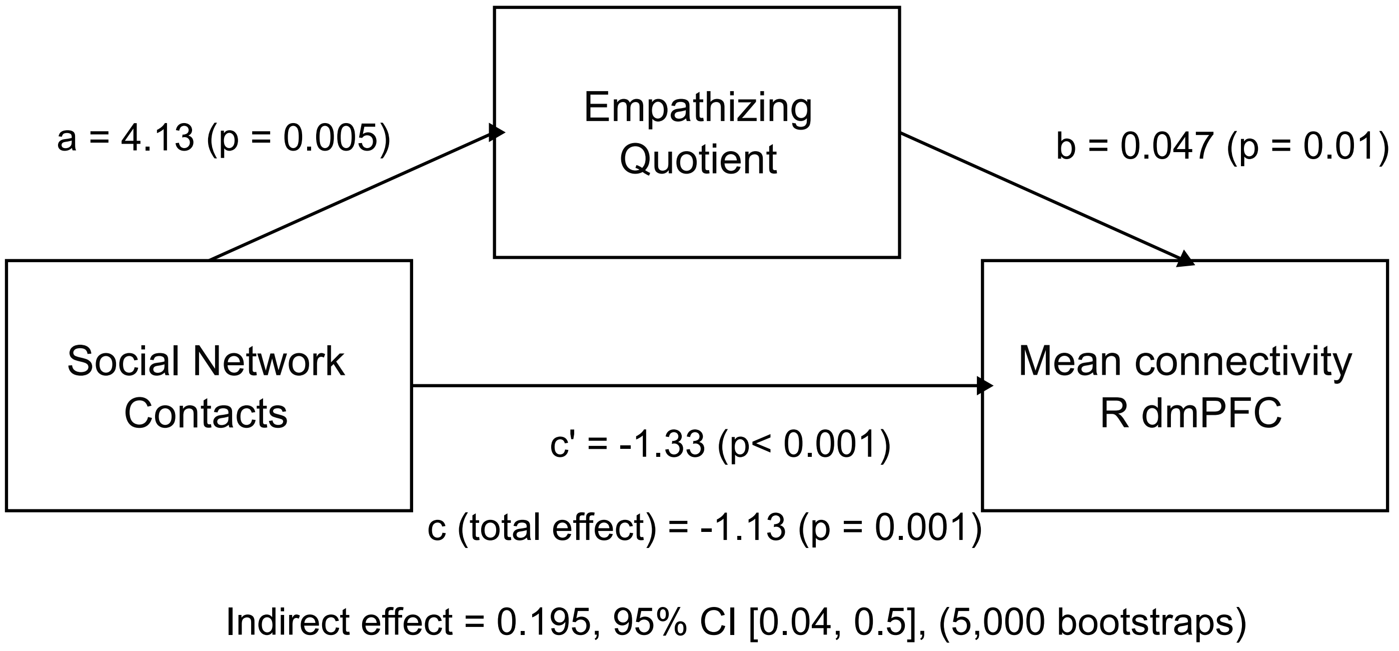


*Legend:* Post-hoc mediation analysis. Unstandardized regression coefficients are shown along with p-values. Mediation analysis was boot-strapped 5,000 times and CIs did not overlap with zero, indicating a significant partial mediation. R dmPFC = right dorso-medial pre-frontal cortex.
